# Genetic Analysis of Deep Phenotyping Projects in Common Disorders

**DOI:** 10.1101/197459

**Authors:** Elliot S. Gershon, Godfrey Pearlson, Matcheri S. Keshavan, Carol Tamminga, Brett Clementz, Peter F. Buckley, Ney Alliey-Rodriguez, Chunyu Liu, John A. Sweeney, Sarah Keedy, Shashwath Meda, Neeraj Tandon, Rebecca Shafee, Jeffrey R. Bishop, Elena I. Ivleva

## Abstract

Several studies of complex psychotic disorders with large numbers of neurobiological phenotypes are currently under way, in living patients and controls, and on assemblies of brain specimens. Genetic analyses of such data typically present challenges, because of the choice of underlying hypotheses on genetic architecture of the studied disorders and phenotypes, large numbers of phenotypes, the appropriate multiple testing corrections, limited numbers of subjects, imputations required on missing phenotypes and genotypes, and the cross-disciplinary nature of the phenotype measures. Advances in genotype and phenotype imputation, and in genome-wide association (GWAS) methods, are useful in dealing with these challenges. As compared with the more traditional single-trait analyses, deep phenotyping with simultaneous genome-wide analyses serves as a discovery tool for previously unsuspected relationships of phenotypic traits with each other, and with specific molecular involvements.

---

The major medical successes of genetic association have been with patient-control comparisons, initially in single-gene diseases and more recently in common (multifactorial/polygenic) diseases and syndromes. In these analyses, disease is often used as a categorical phenotype, present or absent, ignoring the underlying clinical, genetic, and biological complexity of many medical and psychiatric disorders. Recently there have been successful associations with continuous complex phenotypes, such as height, and educational attainment (Rietveld et al., 2013; Wood et al., 2014), and with components of disease, such as blood pressure (Liu et al., 2016; Surendran et al., 2016). In psychiatric disorders, systematic studies of disease aimed at component phenotypes and their biological basis, based on diverse phenotypic measurements, may offer special promise for illuminating their genetics. This view has led to the development of several studies of multiple phenotypes in which many clinical, behavioral, neurophysiological, and neuroanatomic phenotypes, as well as genotypes, are assessed for each studied person in large samples of individuals. We refer to these as deep phenotyping studies; the NIMH-supported Bipolar-Schizophrenia Network on Intermediate Phenotypes (B-SNIP), which has generated this paper, is one such project. The genetic analysis of a deep phenotyping study presents its own challenges, because of the large number of tests, limited numbers of subjects, imputations of phenotypes and genotypes, methodological diversity of phenotype measures, and because of the cross-disciplinary nature and multiple collaborators involved in such studies. The diverse approaches to investigating genotype-phenotype relationships can lead to a confusing array of findings which may often appear contradictory; a comprehensive perspective on the advantages, limitations and tradeoffs involved in the different approaches can provide a clearer perspective for the field.

## A. Genetic architecture of common traits and diseases

Quantitative neurobiological traits related to common neuropsychiatric diseases have become of particular interest since the publications on Research Domain Criteria (RDoCs) (Insel et al., 2010; Insel and Cuthbert, 2009), which are expansions of the endophenotype concept that had been proposed decades earlier by Gottesman (Gottesman and Gould, 2003; Gottesman and Shields, 1973; Gottesman and Shields, 1972). Biological markers, phenotypes, and underlying neurobiological functions related to disease are conceptualized as continuous “domains” that are commonly show varying degrees of abnormality in patients. The implicit theory on the genetic architecture of disease is that there are multiple genetic variants that are correlated with trait markers, and with the right genetic combination the trait markers’ quantitative value crosses a threshold for genetic liability to disease. An alternative architecture would be when, the endo- and/or clinical phenotype appears continuous, but in ill people different genes are operating to produce extreme values, or illness overrides the genes that influence the trait in well people. Yet another complexity of genetic architecture would be when single rare genetic variants can by themselves produce very substantial risk of disease and trait abnormality.

With the development of genetic technologies and analytic methods in the past decade, these and other hypotheses on the roles of genes in trait biomarkers related to disease can be tested directly. In this paper, we discuss genetic analysis of traits that may underlie psychosis syndromes, and detection of genetic architectures of sub-phenotypic traits. We pay particular attention to comprehensive analyses of a wide range of neurobiological phenotypes, and of genomic events associated with these phenotypes (variations in genotype, gene expression, or epigenomic measures), and their relation to individual case/control status.

Among the types of genetic architectures of illness and neurobiological traits, as discussed in the preceding paragraph, there are single gene disorders/traits, quantitative traits with threshold for disease, different genetic events in patients vs. controls, and different genetic events in different ethnic backgrounds. For illness and other traits, a general principle was elucidated by Manolio (Manolio et al., 2009) (Figure 1).

**Figure 1.**
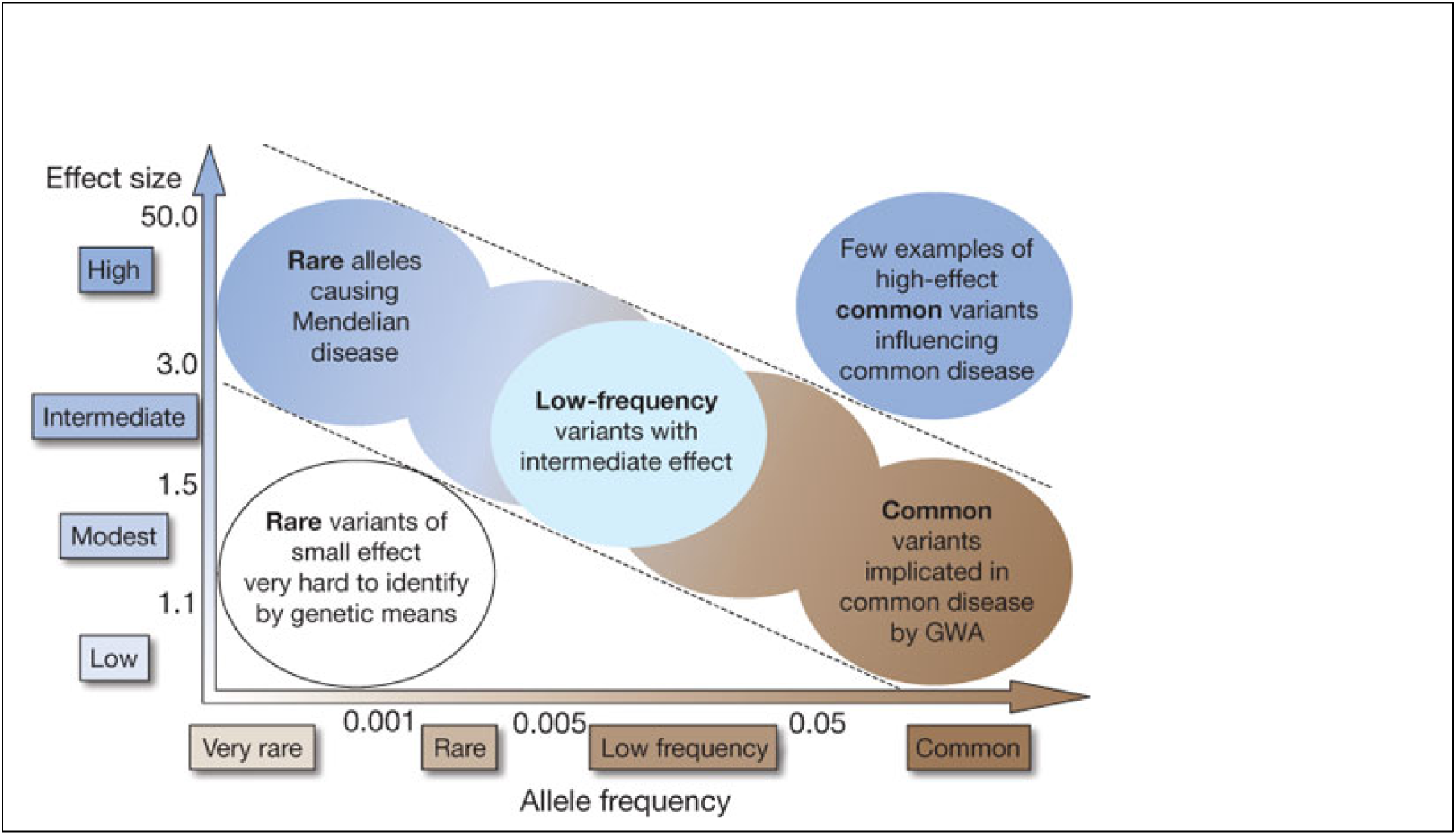
Feasibility of identifying genetic variants by risk allele frequency and strength of genetic effect (odds ratio). From Manolio (Manolio et al., 2009).

With the exception of rare variants with a large effect size (on disease risk), a polygenic model of common diseases, at the lower right of Figure 1, with multiple common genes of weak effect, fits schizophrenia and bipolar disorder, and may be applicable to some marker phenotypes we discuss in this paper.

The term polygenic originated decades before there was a genetic map, to refer to genetic influences that are each too small to identify, but which can have a net cumulative influence on a phenotype. Polygenic inheritance with normally distributed genetic liability was first proposed by Sewall Wright (Wright, 1934), to account for inherited discrete phenotypes. The first applications were to inheritance of the number of digits in guinea pig paws, which could take one of two values. This type of inheritance was recently depicted graphically by Felsenstein (Felsenstein, 2005) (Figure 2). By inspection, the graphic makes it clear that the phenotype as well as the genotype can be continuous, with a threshold for a binary trait (such as illness). For genetic association analysis, there is a statistical appeal in phenotypically continuous traits, because there would be enhanced statistical power to detect genetic associations as compared with the same trait analyzed as a binary phenotype. For many common disorders, including Type 2 diabetes, schizophrenia, and bipolar disorder, there are sub-threshold diagnostic states found in persons at increased risk of illness, and in some family members of patients, suggesting that a quantitative trait analysis of some phenotypes across patients and controls would be appropriate.

At the present time, when there is a dense map of human genetic markers, it is possible to use genotypes to directly assess polygenic risk to a phenotype. In the psychiatric diseases, there is a mixture of common variants with low effect, which can be summarized in a polygenic risk score (PGRS) (Purcell et al., 2009) which has a larger effect than any single variant, but is only currently applicable in Caucasian ancestry persons. The same study showed the schizophrenia PGRS to be applicable to bipolar disorder but not to several medical disorders. The schizophrenia PGRS has also been associated with cognitive function in healthy individuals but not in patients with a history of psychosis (Shafee et al., 2017), suggesting that disease related factors may in some circumstances overwhelm the genetic influences typically influencing a trait in the general population [cf. (Hochberger WC, In Press)]. A mirror image example is found in genetic effects on intellectual ability and disability. The normal range of intelligence is polygenic (Sniekers et al., 2017), but intellectual disability (IQ < 50) is overwhelmingly associated with single rare variants (Gilissen et al., 2014).

**Figure 2.**
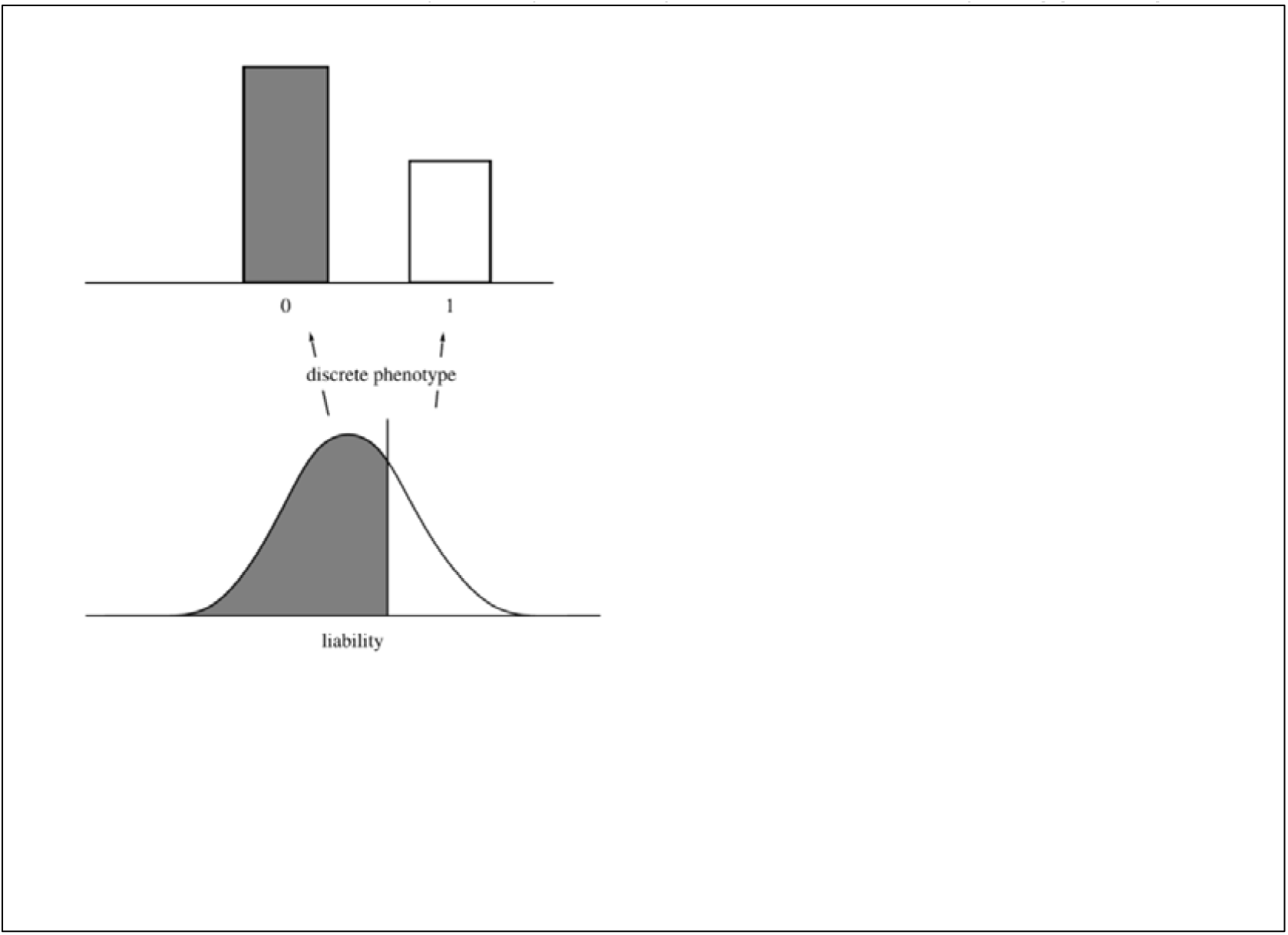
Sewall Wright’s 1934 concept of polygenic variation. “The threshold model of quantitative genetics, showing the continuous distribution of the underlying liability characters, and the resulting discrete distribution of the phenotype.” (Felsenstein, 2005) (see text).

## B. Genome-Wide Association Study (GWAS) of common variants in polygenic diseases or traits

Strictly speaking, the polygenic model does not include variability among the genetic loci, where some loci have more effect than others. But this is the case when genome-wide markers are applied to large disease datasets. Even though many or all genes may contribute to heritability, it is generally agreed that there are “core genes” that are the most interesting to study, such as genes whose SNPs have the largest effect sizes (Boyle et al., 2017).

It is arguable that GWAS is a fishing expedition, so it would be more sensible to test specific molecular hypotheses. Empirically, however, the experience of genetic associations in common disease has generally not supported the candidate gene approach. The GWAS approach has had notable successes, particularly in schizophrenia (Ripke et al., 2014). Unlike fishing the oceans of the Earth, the genome has a finite and encompassable number of genetic locations to study.

GWAS currently refers to association of phenotypes with common single nucleotide polymorphisms (SNPs; minor allele frequency (MAF) >5%). Using well established methods of genotype imputation, which is the process of replacing missing values with estimated data from whole-genome sequences of thousands of people, the entire genome is covered (Howie et al., 2012; Marchini and Howie, 2010). Genotypes of over 4.3 million common SNPs can be imputed from a DNA microarray-based genotyping procedure on ∼500,000 SNPs.

In common diseases, GWASs have mainly studied diagnosis as a categorical (qualitative) phenotype, with relatively few non-diagnostic measures being examined. However, in psychiatric disorders there now exist a number of consortia with extensive genotypic data on hundreds or more phenotypes in thousands of patients and controls. Examples include B-SNIP, which studies multiple categories of neurobiological traits in living patients and controls, the ENIGMA consortium that studies brain imaging, and the Common Mind Consortium which studies autopsy brains. It is in the nature of deep phenotyping studies that the number of individuals studied is relatively small. None of these consortia has the tens of thousands of individuals who have had serum lipids analyzed (Pirruccello and Kathiresan, 2010), because evaluating large numbers of non-diagnostic phenotypes is not part of medical practice in psychiatry. The data are expensive and difficult to collect, and it is a question of research strategy how best to analyze genome-wide data on many phenotypes in limited-size samples.

To improve the likelihood of detecting significant associations, we would offer several principles. First, wherever it is reasonable to do so, it is preferable to study quantitative phenotypic variables in genetic association studies. Whether the quantitation is meaningful should be considered in advance of the genetic analysis, of course, but demonstration of heritability of a quantitative trait can be an outcome of the genetic analysis, and this has bearing on meaningfulness.

Second, it is a statistical fact that a correlation present in a sample is attenuated in a subsample with a smaller range of values, and this applies to phenotype-genotype correlations. The general principle of preferring genetic analysis of samples with the widest variation on quantitative variables was derived long ago for genome-wide linkage studies (Risch and Zhang, 1995), and it applies to association studies as well. Subdividing a sample of individuals into separate groups with different ranges of phenotypic values before doing an integrated GWAS, which can happen when doing separate GWAS on patients and controls, will decrease the likelihood of detection of genetic variations that apply to the entire sample of cases and controls grouped together. There will be instances where a genotypic association is detectable only in one subgroup and not in the collection as a whole. This can often be tested by including appropriate covariates in analysis of an inclusive sample set. Separate analysis of cases and controls can be reasonable as a secondary analysis, such as when it is suspected that illness is masking genetic effects, but there is much to lose by starting with separate analyses.

Third, genetic differences between continental ancestry groups constitute a special case of heterogeneity within a sample. There are methods for analyzing mixed continental ancestry data, with the current preferred method being analysis of the genotype data with Principal Components Analysis (PCA), and adjusting the genotypes by the Principal Components that take up [the most] variance (Price et al., 2006). Meta-analysis of results from the separated groups can also be done. The possibility that there are separate genetic controls of the same phenotype in two different ancestries should be considered only with great caution. Support would be needed for such an interpretation, such as very different allele frequencies at the associated loci.

### a. The multiple testing issue and thresholds for GWAS significance

When imputed genotypes are studied, with imputation applied to the non-genotyped common SNPs, 5E-8 has been suggested by Sham and Purcell (Sham and Purcell, 2014) as a general threshold for significance for a European ancestry sample. This is based on α (false positive rate) of 0.05 for a single test, and a calculation of non-independence (based on linkage disequilibrium) among the 4.3 million imputed genotypes. However, when imputation is applied to more than one continental ancestry group, the recommended significance level for a single GWAS is 1E-8 (Li et al., 2012). We will call this the single-test threshold. The Bonferroni project-wise threshold would be the single test threshold divided by the total number of GWASs in the project. Simulations across all the phenotypes studied can generate an empirical project-wise threshold, at the cost of a great deal of computation.

For deep phenotyping studies, we would add another threshold for significance – within-topic significance. If there are a group of tests (such as morphometric brain measures), they would constitute a within-topic group, and a Bonferroni correction based on group-wise error for that group of tests could also be presented. If there are 100 tests within a group, and the single-GWAS significance threshold is 1E-8, the group-wise Bonferroni threshold would be 1E-10. Results in a deep phenotyping project could be reported as significant for a single test, for a group of tests in a related topic, or for all the tests. Similarly, groups of genes may be considered separately from the whole-genome association study, as in gene-set analysis.

The reason for these several thresholds is to take into account that the number of tests may proliferate endlessly in a study with many complexly interacting phenotypes and data open for public analysis. Each analysis has an expected rate of false positives, and the same genotypes are being permuted, as it were, for each GWAS. The project-wide threshold would apply to all GWASs performed on the sample. If 1000 GWAS analyses are performed, the Bonferroni correction would give a significance threshold of 1E-11, and these would be the strongest results, if statistical power is acceptable (see next section).

The underlying problem for statistical analysis is to estimate the probability of a false positive result in each group of results derived from a single set of genotypic observations. The probability of a false positive in at least one of the phenotypes for which GWAS is performed increases with each GWAS (or gene subset analysis). It is tempting to publish bits of results separately, and to restrict multiple test correction to the genotypes in one GWAS. But this gives a false picture of the sample space, and would add to the unfortunate number of existing false positive GWAS results (Ioannidis, 2007). In our example of 1000 GWASs, 1000 papers could be published with an average of one significant result each, of which only 1 would reach the project-wide significance threshold.

There are alternatives to Bonferroni correction of multiple tests. Bonferroni correction is the upper bound of conservative correction, treating each test as independent, but other less restrictive corrections such as False Discovery Rate (FDR) can be applied. The FDR statistic essentially looks at the distribution of p-values to determine significance, and this can be applied to a data set with a very large number of p-values from multiple tests. Sham and Purcell note that a q-q plot (of negatively ranked log p-values against their null expectations) displays the same information as an FDR, and has the additional advantage that early expansion of negatively ranked p-values is a strong indication of population stratification (Sham and Purcell, 2014). For deep phenotyping, the q-q plots of the multiple GWASs could be combined.

The potential phenotype space is extremely large, given the number of parameters that can be collected in behavior, physiology, chemistry, and anatomy in a deep phenotyping study. As resolution of measurement increases, the number of measurements increases exponentially. This raises the issue of optimizing resolution for association studies, as possible data points extracted from functional and anatomic imaging and electrophysiological data along with clinical data points can easily rise into many thousands with adverse implications for statistical power. In whole-genome genotype-based association analyses, this is compounded by the “baseline” Type I error correction for a single test, 1E-8.

Strategies for constricting the phenotype parameter space may be considered. On an ‘a priori’ basis, one could prioritize a set of phenotypes for analysis, leaving the remaining parameters in one or more sets of variables for exploratory analysis. This would parallel separation of hypothesis-driven vs. exploratory analyses in power analysis for experimental research, and identification of primary vs secondary outcomes in clinical trials. It is also possible to have sample-determined parameters to select phenotypes for a primary analysis, such as effect size, heritability of the phenotype, or robustness of case – control differences.

However, some of these parameters are not determinable without the GWAS analysis. Heritability as determined from correlations with relatives is not well correlated with SNP-determined heritability, and may not be available for all observations in a studied sample. Case-control differences are important for clinical investigation, but genetic association with directly measured biological processes may be indirectly related to case-control differences (as in interaction with other genes or phenotypes), or be of intrinsic interest. With respect to genotype association, the history of whole-genome association analysis in mental disorders has been that prior candidate gene hypotheses from neurobiology and neuropharmacology have generally failed to be supported, with only a few exceptions. In schizophrenia, the important exception is the major histocompatibility complex (MHC) locus region on Chromosome 6, which has been reported to be associated with schizophrenia for decades (Goldin and Gershon, 1983), and another exception is the modest association recently reported with DRD2 (Ripke et al., 2014), the key gene in the dopamine hypothesis of SZ.

Careful consideration needs to be given to these questions of analysis strategy, as decisions made define the number of phenotypes of primary interest and therefore experiment or project-wise Type 1 error prediction for statistical genetic analysis. Of course, ideally this is done as part of study planning as it has implications from a power analysis perspective. But any prospective power analysis in an association study depends on speculation about crucial parameters such as genotype-specific effect sizes, that can only be determined post hoc (see next section)(Mayr S, 2007). In practice, power analyses can change over the course of a large deep phenotyping project, as new results become available, and new measures of phenotypes such as specific aspects of brain anatomy are developed.

While a priori phenotype selection is appealing from a statistical power perspective, we caution that there is an opportunity cost in not casting a wide net. Our current analysis of B-SNIP1 (the initial B-SNIP project with 977 unrelated probands), which is submitted for publication and is available on biorxiv (http://www.biorxiv.org/content/early/2017/08/11/175489) is an instructive example (Alliey-Rodriguez et al.). In B-SNIP1 there are 463 phenotypes that investigators thought could plausibly be hypothesized as having genetic association with illness or with related biological variables. We used 2.16E-11as the Bonferroni project-wide threshold for significance, 1E-8 for the single analysis threshold, and set group thresholds in-between according to the number of tests in the group. After removing linkage disequilibrium (LD) overlapping signals for the same phenotype, there were 34 results that passed the single analysis threshold, and two that passed the project-wide threshold. Only 4 of the 34 p<1E-8 associations were variables that would have met a standard for inclusion in a primary analysis, based on the data having previously been published and/or described as important components of a phenotype. No tested multivariate score met any threshold for significance. The two findings with project-wide significance were not of major prior interest. Using the actual genotype and phenotype data, the power to detect association at a p-value of 2.13E-11 for these two phenotypes was 61% and 62%.

In this instance, at least, there was little to gain by reducing the number of tests. If the 463 variables had been pared to 46 (that is, by a factor of 10), the project-wide significance threshold would have been one order of magnitude changed, that is, changed to 2.13E-10. Only 1 of the 32 results that missed the earlier threshold would have thus become project-wise significant. This result was a second association with one of the two phenotypes that passed the more stringent project-wide threshold for 463 tests. This phenotype and its molecular associations will be of further research interest based on these results, but given the sense of the collaboration it is safe to say the phenotype would not have been included after a paring process prior to the GWAS analysis.

In each analysis of multiple phenotypes in a study like this one, the investigators’ judgment will play the major role in which tests are performed, and the reviewers and scientific community will decide which results are credible. No sensible investigator would decide to do voxel-by-voxel GWAS in a brain imaging study.

Limiting the tests to be performed may not prove to be a long-term executable strategy for controlling the multiple test “burden”. It is in the nature of deep phenotyping studies that the data become public, data-mined, and subject to many genetic analyses and machine-learning, to discover cryptic associations.

### b. More on statistical significance and power

Several accepted conventions should be stated to begin this discussion, despite their familiarity to many readers of this paper. Statistical tests evaluate whether data as observed departs from a null hypothesis. Commonly, the test estimates the probability (p-value) of departure of observed data from [the centrality parameter of] a data distribution under the null hypothesis, such as t-test, chi square, or F distributions. Alpha (α) is the threshold probability value (p-value) of the test for declaration that the data are significantly different from the null hypothesis distribution. This is the “Type I error” (false positive rate) that is acceptable. Beta (β) is the probability of retaining the null hypothesis when it is false (Type 2 error), and 1-β is the power of the test (the probability of rejecting the null hypothesis when it is false).

For a genetic test of association of a single two-allele locus with a binary categorical trait (such as ill or well), power is a function of the number of individuals tested, the effect size of the set of genotypes at the locus (typically represented for a categorical trait by the odds ratio of risk allele frequency of individuals with each trait category), prevalence of category/disease, allele frequencies, and a disequilibrium parameter (which can be ignored for this discussion). For association with a quantitative trait, the effect size would be the proportion of phenotypic variance attributable to the genotypes at the locus. Based on user-supplied values, the Genetic Power Calculator(Purcell et al., 2003) returns the number of cases and controls needed for the desired power to detect an acceptable association.

A very insightful discussion of the imponderability of parameter estimates required for this type of power analysis, which can be termed “a priori”, was published some years ago (Mayr S, 2007). The effect size may not be knowable until the experiment is performed, and the number of cases and controls may not be under the experimenter’s control. Typically, “reasonable” values of the parameter are given in a proposal, and a decision on performing an experiment is made. However, the effect size and power in a particular study, especially one studying novel phenotypes not previously extensively explored in association studies, will only be estimable “post hoc”, once the data are known. “Post hoc” power analyses can be characterized as instruments providing for a critical evaluation of the (often surprisingly large) error probability associated with a false decision in favor of the null hypothesis.

Since statistical power and effect sizes are closely related to sample size, hypotheses that failed to find support in one study can be found to be supported in a larger study. This is not an inconsistency, it is in the nature of statistical tests. Although power to detect is very important for validity of an association, failure to detect does not mean that there is no association. In 2011, Ripke et al. (Ripke et al., 2011) reported “seven genome-wide significant schizophrenia associations (five of which were new) in a two-stage analysis of 51,695 individuals.” Three years later, 152,764 individuals were studied, and there were 108 loci that met conservatively-defined statistical significance (Ripke et al., 2014).

In a 2014 review on statistical power and significance testing in large-scale genetic studies, Sham and Purcell (Sham and Purcell, 2014) demonstrate that “even among association results that reach the genome-wide significance threshold, those obtained from more powerful studies are more likely to represent true findings than those obtained from less powerful studies.” Significance testing in both genome-wide and exome-wide studies must adopt stringent significance thresholds to allow multiple testing, and it is useful only when studies have adequate statistical power, which depends on the characteristics of the phenotype and the putative genetic variant, as well as the study design.”

## C. Other analysis topics

### a. Genotype and phenotype imputation

In almost every human study with deep phenotyping there will be missing observations on a considerable proportion of subjects, which limits the available number of subjects for multiple-phenotype analysis. Many imputation methods have been developed in biomedical statistics to predict the value of missing phenotypes from the existing phenotypes of an individual, based on the relationship of phenotypes to each other in the dataset. Discussion of the different methods and algorithms for imputation of phenotypes is beyond the scope of this paper. We note that it is important, in each instance of imputation of a new class of phenotype, to document the accuracy of imputation. For example, a learning set and a testing set can be created from existing data, and the accuracy can be measured by the correlation between the imputed and actual data in the testing set.

Marchini et al. (Marchini et al., 2007) developed a method for multipoint genome-wide imputation of common genetic polymorphisms which has since been widely adapted. As currently implemented, it imputes several million common polymorphisms from several hundred thousand actual genotypes, with well-demonstrated accuracy. There are practical limits to this and other imputation methods. Missing data may be below the limits of accurate imputation. For example, current genotype imputation is performed by reference to haplotypes of whole-genome sequence data of the 1000 Genomes Project (Abecasis et al., 2010), based on approximately 2500 individuals from 26 populations. This imputation is considered not sufficiently accurate to use with uncommon and rare genotype variants (minor allele frequency < 1%), and such variants should be disregarded in practice, as should loci where the rare variant is the only minor allele. (A method for haplotyping much larger sets of sequence data, that is applicable to rare variants, has been recently been published (Sharp et al., 2016)).

Joint modeling of phenotypes, after GWAS has been performed, can increase discovery by identifying multiple traits that are correlated with the same genotypes. O’Reilly et al (O'Reilly et al., 2012) developed an ordinal regression method (MultiPhen), based on GWAS of multiple traits in the same individuals. The regression performed is phenotype regression upon genotype (rather than the reverse as done in straightforward GWAS). This regression thus tests for the linear combination of phenotypes with maximal correlation with individual SNPs. Phenotypes are thus combined in an unbiased way, although bias may exist in entering the phenotypes into the GWAS analysis.

The covariance among phenotypes, even in unrelated individuals, can be decomposed to a heritable component and a biological component that is not heritable. Dahl et al. (Dahl et al., 2016) in Marchini’s group developed an innovative imputation procedure, PHENIX, which generates a SNP-based heritability of each phenotype, and then for each individual studied imputes the heritable component of that phenotype. They combine this component with a standard imputation of the non-heritable component to predict the phenotype. That is, their method takes into account overall genetic covariance between samples as well as multivariate phenotypic covariance. In that paper, PHENIX generally outperformed the other extant classes of imputation algorithms based solely on phenotypic covariance (that is, had higher correlation with the true simulated or observed values).

### b. Independent Components Analysis (ICA) of Genotypes and Phenotypes, and Genome-Wide Complex Trait Analysis (GCTA)

ICA and GCTA are hypothesis-free statistical methods for analyzing covariation among a single type of variable, and Parallel Independent Component Analysis (Para-ICA) extends ICA to analyze multiple modalities simultaneously (Meda et al., 2012; Pearlson et al., 2015). Para-ICA thus attempts to maximize ‘cost functions’ both within and between complex feature sets (e.g. voxels in MRI scans and gene interactions). Para-ICA simultaneously identifies clusters of associated, likely interacting genes related to phenotypes such as (a) functional brain networks, (b) related structural brain regions, or (c) physiologic processes e.g. EEG patterns or other potential endophenotypes, and shows their relationships (Meda et al., 2010).

Similar to conventional ICA analyses, extracted components or networks are maximally independent within modality and loading coefficients represent variation among individuals. Networks or components extracted from genetic data are groups of interacting SNP loci, contributing with varying degrees to a genetic process affecting a downstream biological function, i.e. linear SNP combinations highly associated with related phenotypes. This reveals novel, biologically relevant associations that might otherwise not be detected due to small effect sizes and modest-sized sample sets. In data with a large number of characteristics, Para-ICA excels at identifying and comparing the most relevant features. Para-ICA also makes assumptions about noise, thus increasing robustness compared to previous methods. Unlike univariate studies [e.g., genome-wide association studies (GWAS)], para-ICA is therefore able to derive clusters of interacting SNPs, each with a weighted contribution toward one or several quantitative traits, and can thus be readily interpreted in the context of illness-associated molecular/biological pathways. The major advantage of this approach in the context of genetics is its power to leverage differential multivariate patterns with minimal loss of statistical power resulting from necessary multiple comparison corrections. Thus it is ideally suited to medium-sized samples in the hundreds to thousands.

Genome-wide complex trait analysis (GCTA) is a variance-covariance analysis of genotypes method for simultaneously assessing the genetic contributions and correlations of each of the genotyped SNPs available to a particular phenotype (Yang et al., 2011). Its most important contributions in psychiatric genetics have been to analyze the SNP-based heritability of individual disorders with GWAS data, and to show the extent of shared heritability of multiple disorders (Lee et al., 2013).

There are relative advantages and limitations of synthetic variables such as PCA components or factor analysis-derived variables combined taxonomic categories, as empirically based phenotypes derived from multiple observations, vs. analyzing phenotypes separately. Synthesized phenotypes have the advantage of potentially better estimating “true scores” of a multivariate phenotype, reducing the number of statistical tests, and accommodating novel case stratification approaches (Clementz et al., 2016), with the potential disadvantage of creating complex variables whose genetics may also be more complex and more difficult to parse than the underlying phenotypes.

### c. Gene networks

There are many ways to categorize genes into groups; a systematic framework has been offered by Li (Li et al., 2015). Networks can be based on *a priori* groupings, constructed from systematic curation of genes that appear together in the scientific literature, or based on correlated expression modules (Zhang and Horvath, 2005) or other functional genomic correlation among genes (Biological level in Table 1). In analysis of genotypic data in human studies, the different networks can constitute sub-sets for the analysis. Methods include formal gene-set analysis, and, when there are a large number of associations, as there now are for schizophrenia, analysis of which annotation categories are more frequent among the associated markers (Ripke et al., 2014; Sniekers et al., 2017).

*A priori* gene-set categories from the multiple types of network in Table 1 tend to be non-mutually-exclusive. Analyses of shared annotations among genes associated with a phenotype can be more appealing. A useful set of tools for such analyses is the DAVID project of the National Institute of Allergy and Infectious Diseases (NIAID) (https://david.ncifcrf.gov/home.jsp). It is beyond the scope of this paper to consider the multiple-testing issues presented by these multiple and often overlapping network-based association analyses. We are aware of no systematic treatment of this question.

**Table 1.**
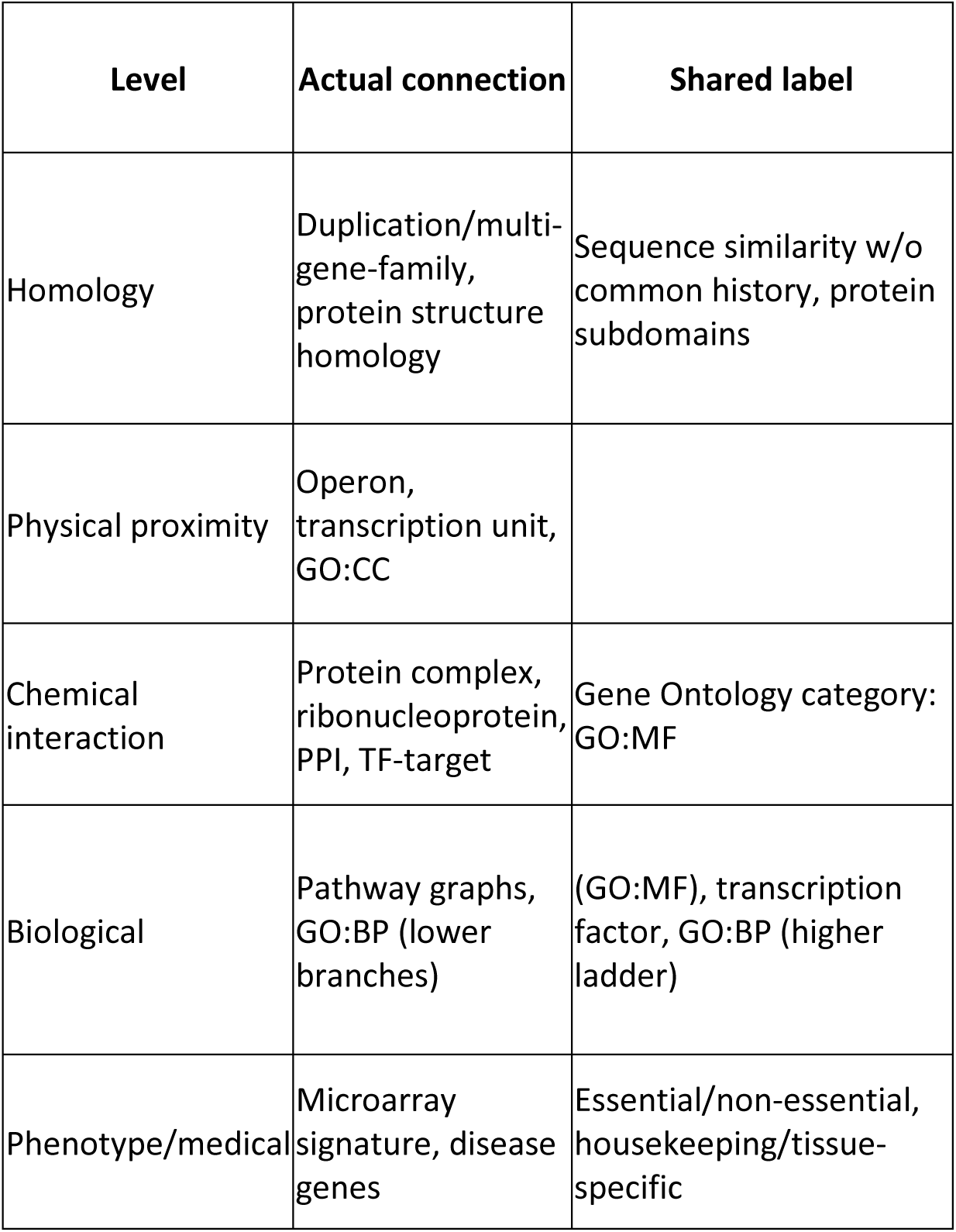
An attempt to assign well known gene-sets under the five-level group headers (homology, physical proximity, chemical interaction, biological, phenotypic/medical) and two types of links among genes in a gene-set (actual connection and shared label). From Li (Li et al., 2015).

| Level | Actual connection | Shared label |
| --- | --- | --- |
| Homology | Duplication/multi-gene-family, protein structure homology | Sequence similarity w/o common history, protein subdomains |
| Physical proximity | Operon, transcription unit, GO:CC |  |
| Chemical interaction | Protein complex, ribonucleoprotein, PPI, TF-target | Gene Ontology category: GO:MF |
| Biological | Pathway graphs, GO:BP (lower branches) | (GO:MF), transcription factor, GO:BP (higher ladder) |
| Phenotype/medical | Microarray signature, disease genes | Essential/non-essential, housekeeping/tissue-specific |

### d. Functional Genomics and Epigenomics

An “omics” suffix is used in biology when the complete sample space of a type of variable can be examined. Thus the human genome is the germ line plus mitochondrial DNA sequence of an individual, and the epigenome is gene expression (the transcriptome) in specific tissues, and all the known classes of event that regulate gene expression. These include DNA methylation, histone modifications, other mechanisms regulating chromatin opening, many long non-coding RNAs, and environment. These events are frequently developmental stage and organ-specific, so there is a probability of some disconnects in studying peripheral epigenomic events, such as in blood element epigenomics, and applying results to the state of the various cells and processes in the brain. Any of these “omics” data, and combinations of them, can be correlated with phenotypes of interest in disease.

For example, differences between patients and controls in gene expression in specific brain regions in autopsy brain specimens have yielded interesting disease associations and associations with specific genes and networks (Cusanovich et al., 2016; Gamazon et al., 2013; Liu, 2011). Sets of SNPs from these genes or networks can be used in genetic association analysis. To a great extent, however, the genomic/epigenomic analyses are a separate set of analyses from genetic (genotypic) association analyses, apparently because the control of functional genomic events can be very complex and not directly correlated with single genotypes. The genes implicated in functional genomic analyses of a phenotype may be different from those implicated by disease association with genotypes.

### e. Rare single-nucleotide variants (SNVs) and copy number variants (CNVs)

Several known structural genomic variants (CNVs, sub-chromosomal deletions. duplications, and translocations) are rare, but when they occur in an individual there is greatly increased risk of multiple mental disorders and intellectual disability (Malhotra and Sebat, 2012); this class of events is included in Figure 1. Persons with these events included in a GWAS along with persons with the “usual” polygenic type of SZ or BD might lead to distortions in genetic and phenotypic analyses. Special collection efforts to assemble samples of patients with the known illness-related CNVs and recurrent rare SNVs may generate enough numbers for statistically valid deep phenotyping investigation, as in the International Consortium on Brain and Behavior in 22q11.2 Deletion Syndrome study of sub-threshold psychosis in that syndrome (Weisman et al., 2017).

## D. Discussion: The value of multiple phenotypes for biologic discovery

Once there is a database with uniform genotyping and genetic analysis on a large number of phenotypes, the advantages of analyzing multiple phenotypes become apparent. Phenotypes related to another phenotype of interest can boost power to detect new associations, allow measuring heritable covariance between traits, and potentially to make causal inferences between traits (Dahl et al., 2016). As compared with the more traditional single-trait analyses, deep phenotyping with simultaneous genome-wide analyses serves as a discovery tool for previously unsuspected relationships of phenotypic traits with each other, and with shared molecular events. Pleiotropy, the spectrum of phenotypes associated with a specific genotype, can also be discovered from comprehensive analyses. Unsuspected instances of phenotypes sharing common genes may come up, as in the discovery of association of an NRXN1 SNP with a ventricular volume and with age of onset of illness, in our own B-SNIP study (Alliey-Rodriguez et al.). Lastly, a comprehensive perspective on all the tested variables in a deep-phenotyping project, and appropriate correction for multiple testing in each publication, may help investigators avoid unreproducible results, which have been unfortunately frequent in the genetics of common diseases (Ioannidis, 2007).

## E. Acknowledgements

Acknowledgements: David Glahn provided valuable advice and discussion on this manuscript.

Grant support: ESG: NIMH MH103368, GP: NIMH MH077945, CT: NIMH MH077851, MSK: NIMH MH078113, BC: NIMH MH103366, JAS: NIMH MH077862.

